# Modelling of pH-dependence to develop a strategy for stabilising mAbs at acidic steps in production

**DOI:** 10.1101/711416

**Authors:** Max Hebditch, Ryan Kean, Jim Warwicker

## Abstract

Engineered proteins are increasingly being required to function or pass through environmental stresses for which the underlying protein has not evolved. A major example in health are antibody therapeutics, where a low pH step is used for purification and viral clearance. In order to develop a computational model for analysis of pH-stability, predictions are compared with experimental data for the relative pH-sensitivities of antibody domains. The model is then applied to proteases that have evolved to be functional in an acid environment, showing a clear signature for low pH-dependence of stability in the neutral to acidic pH region, largely through reduction of saltbridges. Interestingly, an extensively acidic protein surface can maintain contribution to structural stabilisation at acidic pH through replacement of basic sidechains with polar, hydrogen-bonding groups. These observations form a design principle for engineering acid-stable proteins.

## Introduction

The promise of therapeutic antibodies relies on the ability of the pharmaceutical industry to develop large scale manufacturing processes that can produce safe, economic and reproducible formulations. However, these biopharmaceutical production processes are limited by non-specific interactions between the proteins, which can lead to reversible or irreversible association. At best, these association potentials can lead to batch to batch inconsistencies, but more worryingly they can lead to the abandonment of promising therapeutic leads due to an inability to produce a stable formulation, the loss of target specificity or the development of immunogenicity amongst other issues. One of the major driving forces for reversible association is the solution conditions used to purify monoclonal antibodies, in particular the requirement for low pH solution steps. Currently, most therapeutic mAbs are produced using a similar pipeline. Antibodies are normally expressed using mammalian expression systems, usually CHO cells (Birch and Racher, 2006), once secreted the antibodies are then purified using chromatographic and membrane filtration steps to remove impurities. Protein A chromatography was one of the first protein purification processes developed (Duhamel *et al*., 1979), and is still the state of the art for the manufacture of biotherapeutic antibodies (Gronemeyer *et al*., 2014; Ulmer *et al*., 2019) as it is highly effective, resulting in > 98% purity in a single step (Shukla, Abhinav A and Thömmes, Jörg, 2010). It is also highly stable, retaining specificity over multiple uses (Brorson *et al*., 2003). However, the use of protein A necessitates the use of a low pH buffer to elute the protein from the column, and this acid titration can induce antibody aggregation leading to protein formulation issues (Shukla *et al*., 2007; Liu *et al*., 2010; Shukla, Abhinav A and Thömmes, Jörg, 2010). It has also been suggested that the column itself may contribute to aggregation (Mazzer *et al*., 2015). Following protein A purification, antibodies expressed from mammalian sources also require a low pH hold for viral inactivation which can lead to further aggregation (Shukla *et al*., 2007). As well as aggregation, it has been observed that exposure to acidic pH can cause polyreactivity, which may be part of the natural immune response, but is an issue for developing specifically targeted therapeutics (McMahon and O’Kennedy, 2000; Dimitrov *et al*., 2010).

In light of this, attempts have been made to modify protein A to allow the use of less acidic buffers (Ghose *et al*., 2005; Gülich, Susanne and Uhlén, Mathias and Hober, Sophia, 2000; Hober *et al*., 2007), to optimise the resin (Liu *et al*., 2015) and washes (Shukla and Hinckley, 2008), or develop alternative chromatographic processes (Hanke and Ottens, 2014). The development of alternative purification techniques such as pH conductivity (Chen *et al*., 2019) and engineered proteins with a calcium dependent binding for IgG (Kanje *et al*., 2018) has also been discussed. These techniques however do not remove the need for a low pH hold for viral inactivation, instead alternative techniques have been suggested (Burnouf and Radosevich, 2003; Klutz *et al*., 2016). The use of excipients like arginine to help minimise damage from low pH treatments has also been reported (Yamasaki *et al*., 2008).

Due to the requirements of protein A chromatography and viral inactivation, many reports have investigated antibody pH sensitivity and aggregation pathways (Vázquez-Rey and Lang, 2011; Brummitt *et al*., 2011a,b; Arosio *et al*., 2013; Liu *et al*., 2016; Imamura and Honda, 2016; Skamris *et al*., 2016; Imamura *et al*., 2017). To better understand the cause of acid induced protein aggregation, research has focussed on understanding if aggregation is driven by specific domains within the antibody. The IgG is routinely divided into its two functional substructures, the Fab which contains the antigen binding region, and the Fc plays an important role in immunological signalling and activation (Figure 1). These substructures can be further subdivided into individual domains. The Fab contains the variable heavy (VH) and variable light (VL) domains, named for their role in containing the hypervariable complementarity determining regions which bind the antigen, and the constant heavy (CH1) and constant light (CL) named for their conserved sequence. The Fc contains two further conserved heavy chain domains: CH2 and CH3.

**Figure 1:**
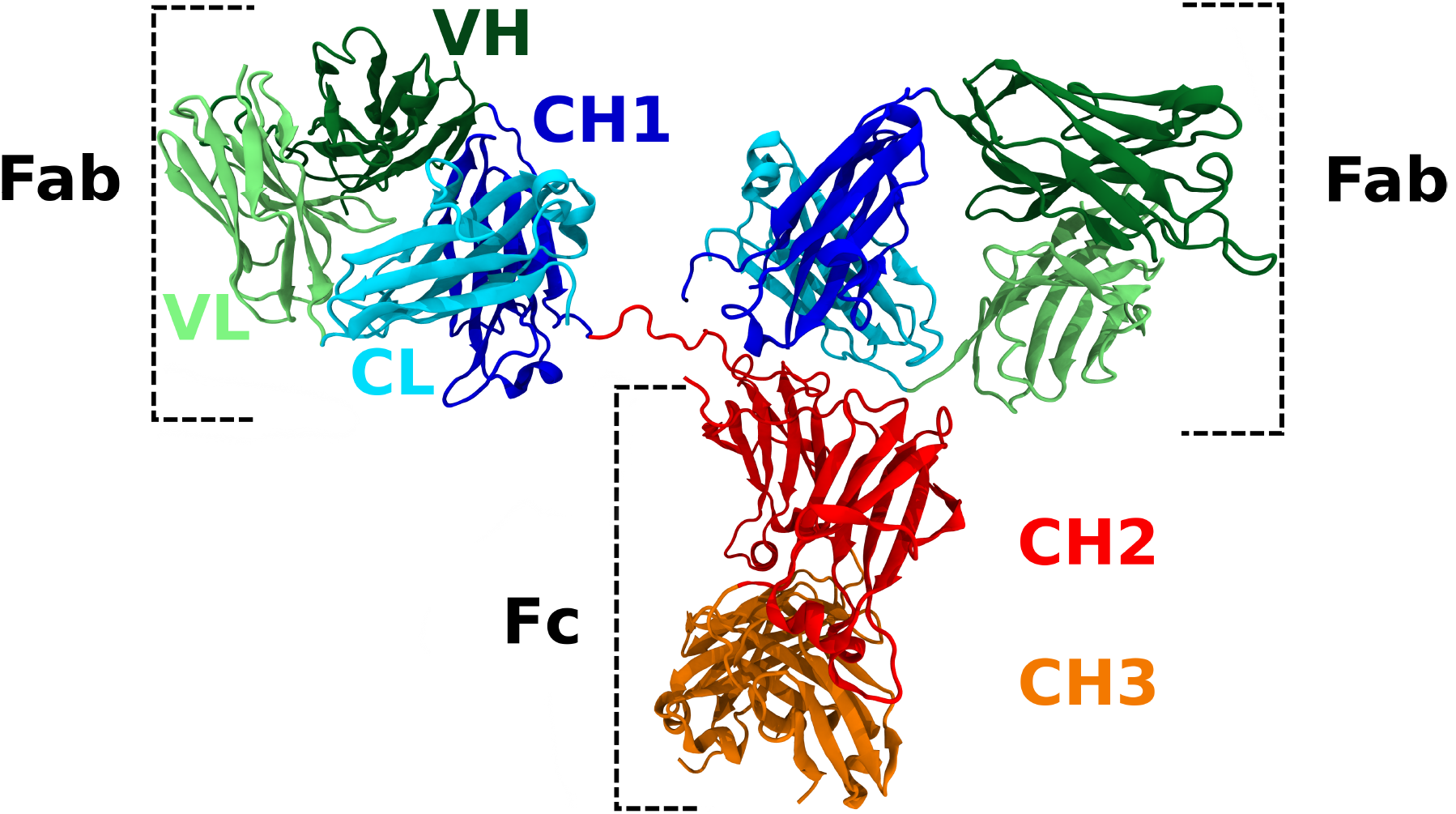
Example schematic of a IgG (1HZH, Saphire *et al*. 2001). Each IgG is comprised of two heavy chains (each containing a VH, CH1, CH2 and CH3) and two light chains (each containing a VL and CL).

The importance of electrostatics in stabilising proteins has been well studied in the literature, however, the contribution of ionisable group interactions to the stability of the folded state will differ between the charge environment that proteins have evolved for, in the case of antibodies physiological pH, and the low pH required for protein A chromatography and viral inactivation. A better understanding of the different role electrostatics play in stabilising the antibody at low and neutral pH is therefore important in order to develop more stable antibody formulations. In this work, we use Debye-Hückel calculations (Warwicker, 1999) to study the contribution of the ionisable group interactions to the folded state stability for the IgG, the Fab, Fc and all of the constituent domains. By studying the predicted response of the individual domains to low pH exposure, we attempt to determine which domains may be most sensitive to acid titratation and thus drive low pH aggregation.

Previous work from our group has studied the difference in sequence and structure of the four Fab domains (Hebditch *et al*., 2017). We determined that the CH1 domain has an unusual sequence composition described as being intrinsically disordered like, appearing to have little charge-charge stabilisation, and may instead be stabilised by its interaction with the CL domain. In this work, we report that the CH1 domain appears to be the least destabilised by acid titration, potentially due to its IDP-like characteristics, but more importantly our calculations suggest that the CH2 domain is the most destabilised at low pH, due to a large loss of ionisable interactions which are stabilising at neutral pH, but destabilising at low pH. This observation may provide insight for developing IgG therapeutics which will be resistant to aggregation in the low pH environment required as part of the industrial production of therapeutic mAbs. Through comparison with proteins that have evolved for functioning at low pH, we make suggestions for engineering strategies that could aid IgG domain stability in acidic conditions.

## Method

### IgG domain dataset acquisition

Structures for the Fab and Fc datasets were obtained from the protein data bank (PDB) (Berman *et al*., 2007). The Fab domains were processed as discussed in Hebditch *et al*. (2017), in brief, the PDB was queried for the term “Fab”, and following a number of quality control steps, the resulting structures were analysed for conserved interdomain sequence motif between the variable and constant domains and then cleaved, resulting in 333 unique Fabs and therefore 333 of each of the four Fab domains: VH, VL, CH, CL. Combined these domains also yield 333 heavy and light Fab chains. The Fc domains were obtained in a similar manner. The PDB was queried for the term “FC IgG1” which, excluding full length IgGs, returned 45 X-ray structures. These 45 chains were then aligned using Clustal Omega (Sievers *et al*., 2011) and conserved motifs at the N and C terminus were identified for CH2 and CH3. For CH2 domains, the N’ motif was ‘PSVF’ and the C’ motif was ‘SK[AT]K’. For CH3 domains, the N’ motif was ‘GQPRE’ and the C’ motif was ‘LSL’. Retaining only structures with unique sequences resulted in 12 Fc PDBs and thus 12 CH2 and CH3 domain structures.

### Acid environment proteins acquisition

A dataset of peptidases active at acidic pH was obtained from the MEROPS database (Rawlings *et al*., 2011). The MEROPS database divides peptidases into families of homologous proteins. MEROPS A1 is a family of 27 endopeptidases with an aspartic active site, and are generally evolved to be most active in acidic environments. Structures were obtained from the PDB (Berman *et al*., 2007) for an acid-stable aspartic acid protease (pepsin, 3utl Bailey et al. 2012), an acid-stable xylanase (1bk1, Fushinobu et al. 1998), and an acid protease that has not evolved to be acid-stable (the BACE2 beta secretase 3zkq, Banner et al. 2013).

### Electrostatic calculations

To assess the stability of the various IgG domains and acid environment proteins, we used electrostatics software previously described (Warwicker, 1999; Moutevelis and Warwicker, 2004; Warwicker, 2004) and also available for free as a web application (Hebditch and Warwicker, 2019). This software uses a Debye-Hückel calculation to calculate the pKas and thus the pH-dependent contribution to folded state stability. Using a Monte Carlo sampling of protonation we compare the stability of the unfolded state, where it assumed there are no interactions between ionisable groups, and the folded state. The folded state energy is defined in terms of kJ per amino acid to ensure a normalisation allowing direct comparison of different sized proteins and protein fragments. We used a continuum medium with a uniform relative dielectric of 78.4 (representing water) and an ionic strength of 0.15M.

## Results and Discussion

### Calculated pH-dependence of stability correlates with experiment

The Debye-Hückel approach calculates the electrostatic contribution to folded state stability from pH 1 to 14. Although a protein may not be stable at these extreme pH values, we use them (without accounting for any conformational change) to traverse the extent of ionisation states for the most important ionisable groups. For each pH value, a negative values indicate that the electrostatic contribution (from interactions between ionisable groups) will favour the formation of the folded state. Positive values suggest that the protein is not stably folded due to electrostatic forces alone, and will require an energetic input, or constraint, to maintain the folded state of the structure. We generally note a similar inverse bell distribution for most proteins, structure or domain interrogated, with each exhibiting the most electrostatic stability, evident from the most negative kJ/aa values, between pH 5 to 9, which is to be expected as IgGs have evolved to be stable at physiological pH (Figure 2). At the more extreme pH values, less than pH 4 where the acidic residues are uncharged, and greater than pH 10 where the basic residues are uncharged, we observe positive kJ/aa values indicating that the proteins are less stable the further they are from physiological pH, where either the acidic or basic residues are uncharged and thus cannot contribute to the electrostatic stability of the folded state through ionic interactions.

**Figure 2:**
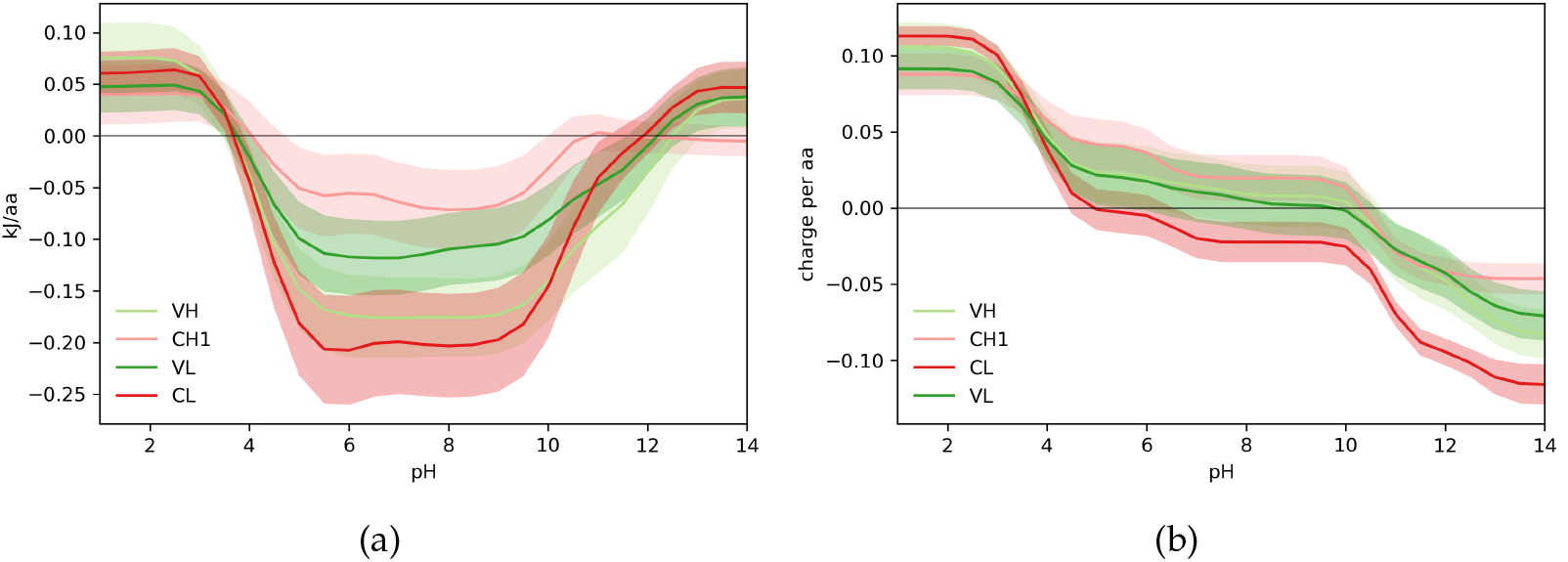
Calculated pH-dependence of folded state energy and charge for Fab domains. (a) The pH-dependent component of folded state energy is plotted against pH, showing average and 1 standard deviation spread for each domain type in the Fab dataset. (b) Predicted pH-titrated charge.

The Fab substructure consists of four domains which appear to have different fold stability profiles. Of the four Fab domains, our calculations suggest that at physiological pH the CL domain has the greatest contribution from electrostatic forces to folded state stability and is therefore the most stable, followed by the VH and VL with CH1 exhibiting the lowest electrostatic stability (Figure 2a). At low pH, the four domains exhibit similarly destabilising contributions from ionisation group interactions. As CH1 has the lowest loss of stability upon transition from neutral to acidic pH, we predict that the CH1 domain is the least destabilised by low pH conditions. CL in comparison loses a large degree of stabilising electrostatic force and is therefore more sensitive to changes in pH.

In terms of charge at neutral pH, the CL domain is calculated to have a negative charge, with the other three domains predicted to be positively charged. Of note, the VH and VL domains appear to exhibit very similar charge, but the CH1 and CL appear to have a difference in charge of around 0.04e per aa at physiological pH and changes in the charge for CH1 and CL appear to be closely related (Figure 2b).

The difference in ionisation energy and charge suggests that within the Fab, the CH1 domain appears to exhibit surprising stability when transitioning from neutral to low pH, and there also appears to be a potentially compensatory relationship with the CL domain. In previous work from our group (Hebditch *et al*., 2017), we noted that the CH1 domain of the Fab sampled an amino acid composition subset and as a result was enriched for certain amino acids (P, S, T, V) and depleted for others (D, E, I, F, Q, R, Y). We described the CH1 domain as being IDP-like in the absence of the CL domain and this is supported by experimental work in the literature that notes that the CH1 domain is not stable in isolation (Vanhove *et al*., 2001; Röthlisberger *et al*., 2005), but exhibited a very stable interaction with the CL domain. Similarly, *γ*B-crystallins are highly soluble proteins with two domains that have similar sequence and structure but fold independently, and by different pathways. The C-terminal domain alone is unstable under acidic conditions but exhibits a high degree of stability in acid environments when complexed with its binding partner the N-terminal domain (Mayr *et al*., 1997). This suggests that acid instability in one binding partner can be abrogated through interaction with an acid-stable partner and the instability of the CH1 domain may therefore be rescued by the high stability of the CL domain. Interestingly, results from the literature suggest that engineering the CH1 domain to contain more disulphide bonds can increase the thermal stability of the Fab (Peters *et al*., 2012). The interactions between the CH1 and CL and the VH and VL appear to be important, and may rescue the less stable CH1 domain, but there appears to be little stabilisation between the CH-CL and VH-VL dimers (Röthlisberger *et al*., 2005), although the CH-CL dimer may play a role in desolvating hydrophobic moieties on the VH-VL dimer that are exposed when the VH and VL are expressed as a scFv (Nieba *et al*., 1997).

After investigating the Fab, we then moved on to compare the Fab as a whole to the CH2 and CH3 domains of the Fc, since there are pH-dependent experimental data with which to compare for these constituents (Figure 3). For example, results in the literature suggest that the Fab has a lower melting temperature than the Fc (Welfle *et al*., 1999; Vermeer and Norde, 2000; Vermeer *et al*., 2000), however the Fc is believed to unfold faster than the Fab in the presence of chemical denaturants (Rowe, 1976), and the Fc is more sensitive than the Fab to aggregation at low pH (Liu *et al*., 2016; Vermeer and Norde, 2000). In terms of ionisable contributions to the folded state stability (Figure 3a), the CH2 domain appears to be more stable than the CH3 domain at neutral pH, but less stable at low pH. The Fab substructure lies between the CH2 and CH3 domains in terms of stability at both neutral and low pH. Notably, out of all of the IgG domains, the CH2 domain appears to be the domain most stabilised by electrostatic contributions at physiological pH, but is amongst the most destabilised at low pH. This is to be expected as proteins with a greater degree of charge stabilisation at neutral pH will be disproportionately affected once these charge groups are neutralised, and therefore proteins that are highly stabilised at neutral pH are more likely to show instability through acid pH titration. From our calculations, the CH2 domain also exhibits the greatest magnitude of charge, as well as the most positive charge at low pH, and is amongst the most charged at physiological pH which may explain both the high level of charge-charge stability at neutral pH, but also the relative instability at low pH (Figure 3b).

**Figure 3:**
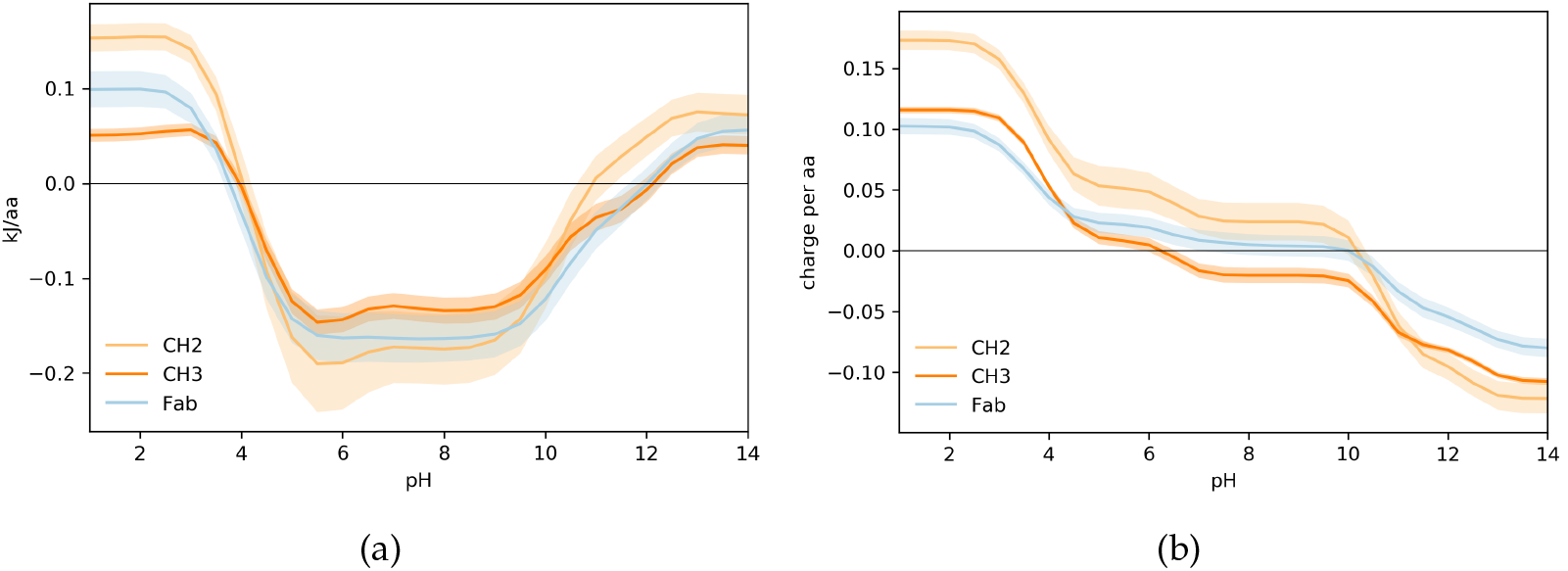
Calculated pH-dependence of folded state energy and charge for Fab and Fc domains. (a) The pH-dependent of folded state energy shown for Fab compared with Fc domains. (b) Predicted pH-titrated charge.

Our understanding is that the pH instability of the CH2 domain is in part related to its high stability at neutral pH, and the importance of the CH2 domain in acid aggregation has also been observed experimentally. The role of the CH2 domain for driving IgG aggregation in an acidic environment appears to be linked to the glycosylation state of the CH2 domain, with aglycosylated mAbs demonstrating a lower Tm (Hari *et al*., 2010; Liu *et al*., 2016; Latypov *et al*., 2012) and fragmentation of the IgG at pH 4 also appears to occur in the CH2 domain (Gaza-Bulseco and Liu, 2008). Also upon exposure to low pH, the CH2 appears to undergo a structural transformation and once renatured by neutral pH titration, leads to an increased thermal stability of 12 – 15°C for the whole IgG (Martsev *et al*., 1994), however this conformation change appears to have little effect on the structure of the IgG overall (Ejima *et al*., 2007). Subsequent work from the same group suggests that before acid treatment, the CH2 exhibits stronger interactions with the CH3 domain, but upon renaturation, the CH2 instead forms stronger interactions with the CH1 domain (Martsev *et al*., 1995). Lastly, experimental determination of the conformational stability of isolated IgG constant domains using circular dichroism suggests that CH2 and and CH3 domains are less stable than the CH1-CL dimer at pH 2 (Yageta *et al*., 2015). Our work suggests that of the Fc domains, both structures exhibit a relative sensitivity to acid pH, but as CH2 exhibits a more destabilising energy in the acidic regime we suggest CH2 will be more acid sensitive.

Thus, to alleviate the relative instability of the CH2 domain in the low pH region, it is not as simple as mutating charge interactions, as stability would then be compromised across a range of pH conditions. Instead, the interplay between the contribution from ionisable group interaction needs to be considered against other stabilising contributions to the folded state. To better understand the importance of the electrostatic stability contribution, and the differences between the apparent relative acid stability of the CH1 and the instability of the CH2, we investigated the electrostatic contribution to stability at low pH for proteins that have evolved for stability at low pH.

### Stomach resident proteases have reduced ionisable charge pair interactions

In order to contextualise the apparent pH instability of the IgG CH2 domain, and relative stability of the CH1 domain, we compared both IgG domains to proteins that have evolved to low pH environments. In Figure 4 we compare the CH1 and CH2 to the Merops A1 family of stomach resident aspartic proteases. The IDP-like CH1 and the acid stomach proteins appear to be less destabilised by the transition to the low pH regime, and are thus less likely to unfold. In comparison, the CH2 domain, which is believed to be acid sensitive but stable at physiologically pH, has a markedly different charge contribution profile to the acid-stable proteins. The CH1 domains and the stomach proteases have reduced ionisable group interactions compared to the CH2 domains, and therefore have less stabilisation to lose upon D/E neutralisation at the low pH found in the stomach (pH 1.5–3.5).

**Figure 4:**
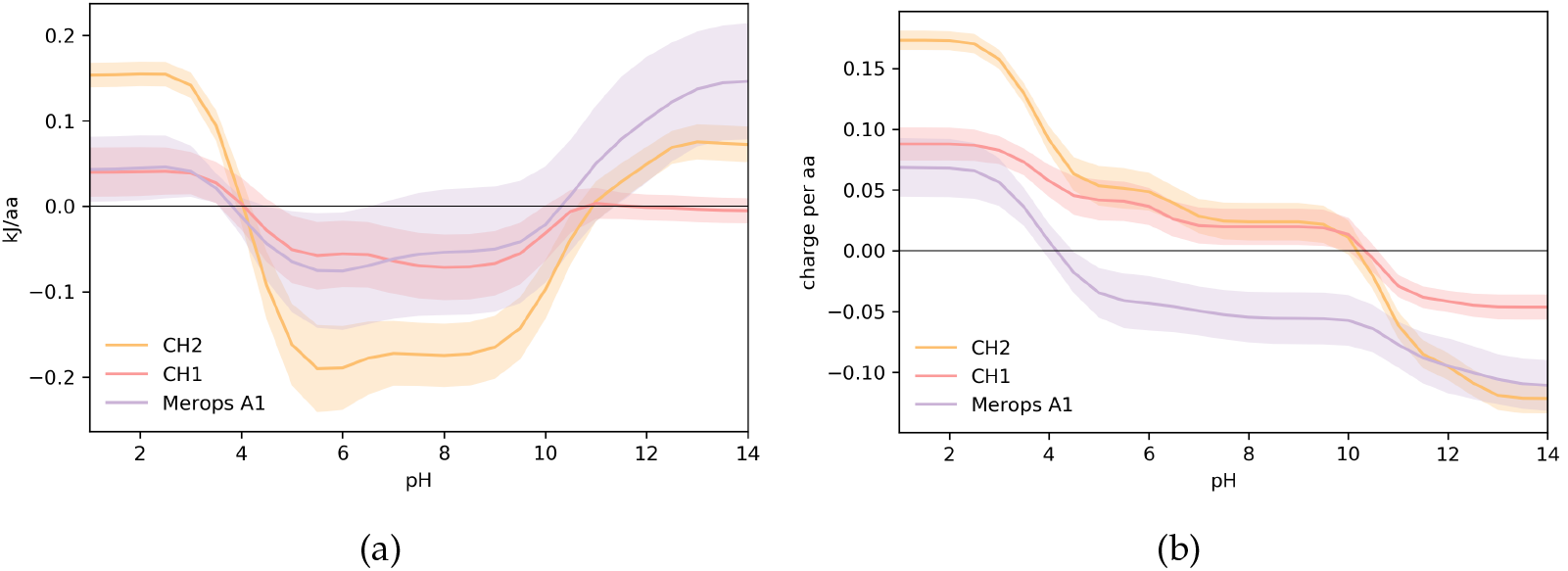
Comparison of calculated pH-dependence for Fc domains and acid-stable proteases. (a) pH-dependence of folded state energy. (b) pH-titrated charge.

Next, we calculated the ionisable energy contribution to folded state stability for a pepsin, a specific acid-stable protease and BACE2, a (human) homologue of pepsin that is not stomach resident. For comparison we also calculated the energy and charge profiles for a industrially important xylanase C (XynC) enzyme from the acidophile *Aspergillus kawachii* (Figure 5) which has an optimum pH of 2 for catalytic activity but is stable at pH 1, and in comparison to mesophillic xylanase C proteins, lacks the characteristic Ser/Thr surface in favour of expressing a negative surface with anisotropically distributed acidic residues (Fushinobu *et al*., 1998).

**Figure 5:**
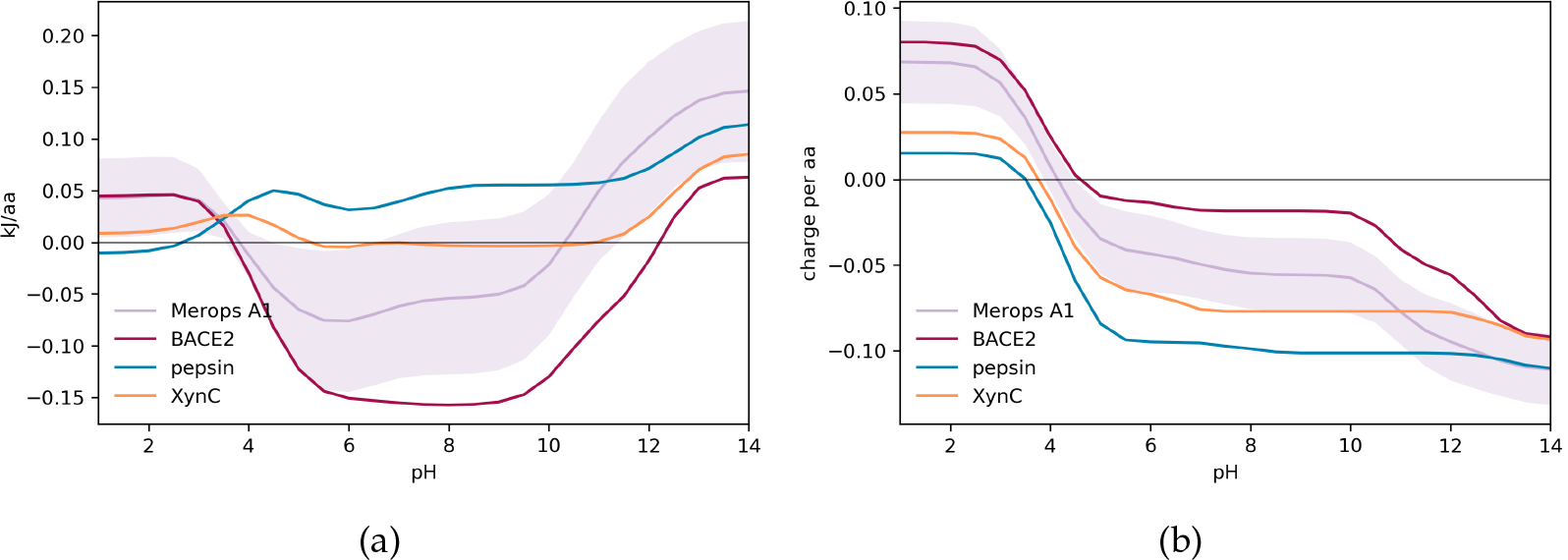
Comparison of calculated pH-dependence for proteins with differing acid-stability. (a) pH-dependence of folded state. (b) pH-titrated charge.

BACE2 is predicted to have a larger contribution from ionisable group interactions, and thus stability, at neutral pH, which is lost at acidic pH (Figure 5a). By contrast, the pH-dependence for pepsin and XynC are much flatter so that although the ionisable group interactions do not contribute greatly to fold stability at neutral pH, they do not lead to acid instability. In terms of charge (Figure 5b), pepsin and XynC are clearly highly acidic proteins. Charged protein surfaces are maintained in acid-stable proteins, but they are not necessarily used extensively in the formation of salt-bridges. Thus, although pH-dependence is flattened, consistent with evolution for an acid environment, the question remains of how charged group interactions are arranged so as to maintain stability at neutral pH.

### Pepsin comparison with BACE2 homologue reveals repurposing of acidic sidechains from salt bridges to hydrogen-bonding interactions

In a closer look at how charge interactions are used for folded state stability at neutral and acidic pHs, specific examples are identified graphically for pepsin and its BACE2 homologue. Figure 6 shows two examples where basic amino acids in BACE2 have been replaced by amino acids with polar sidechains in pepsin. In each case, interaction with an acidic sidechain is maintained, but rather than positive-negative charge pairing, this is now hydrogen-bonding. We infer that at the acidic pH of the stomach, where a positive charge would be left unpaired as a neighbouring acidic group protonates and neutralises, a polar but uncharged sidechain allows for formation of a hydrogen-bond network that can adapt and be maintained through protonation of the neighbouring acidic group.

**Figure 6:**
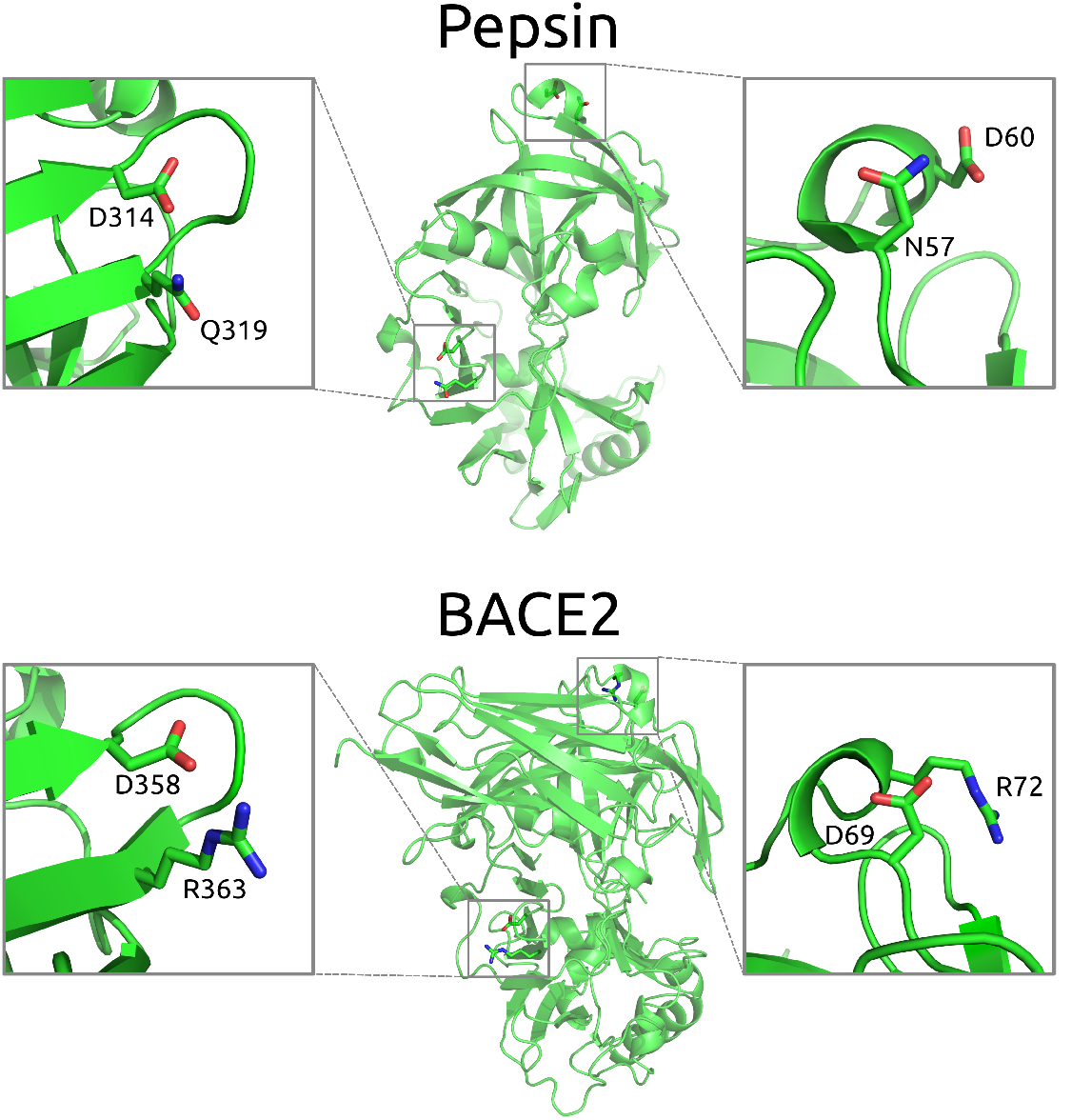
Comparison of acidic group interactions in equivalent regions of the homologues pepsin and BACE2.

In the case of the D358-R363 interaction in BACE2, a glutamine replaces the arginine in the D314-Q319 equivalent pairing for pepsin. In both BACE2 and pepsin, the aspartic acid sidechains also make hydrogen bonds to mainchain groups, presumably contributing to overall fold stability, at least locally in the loop region displayed. Also shown in Figure 6, the D69-R72 charge pair in BACE2 is replaced by the D60-N57 hydrogen bond pair in pepsin. In this case, although structurally equivalent, the relative locations are inverted between the two enzymes.

In general, it appears that pepsin can maintain a stabilising network of negative charges through swapping in hydrogen-bond interactions for charge pair and salt-bridge interactions, giving a basis for redesign of surface charges in domains that are required to be stable at low pH, but have not evolved with that restraint.

In order to show calculations of pH-dependent stability that are accessible with the heat map plots at www.protein-sol.manchester.ac.uk (Hebditch and Warwicker, 2019), the Debye-Hückel (DH) method was used. Previous work shows that this accounts reasonably well for interactions between ionisable groups that are on the protein surface (Warwicker, 1999), whereas buried and partially buried groups are better analysed with a combined Finite Difference Poisson-Boltzmann (FDPB) and DH methodology, termed FDDH (Warwicker, 2004). The FDDH method includes better estimation of hydrogen-bonding energetics to ionisable groups in less solvent accessible regions, generally the type of interactions that we have identified in pepsin (Figure 6). Since FDDH calculations are more computationally expensive than DH, they are not currently available on the protein-sol web server, but local computations for human pepsin and BACE2 pH-dependence have been made. For most proteins, DH and FDDH calculations give (qualitatively at least) a similar predicted pH-dependence of folded state stability, since buried ionisable group interactions do not play the major role in establishing this dependence. This similarity is also the case for BACE2 (Supp Fig 1). However, the DH (Figure 5) and FDDH curves (Figure 7) are quite different in two notable ways. First, there is a steep pH-dependence between pH 4 and alkaline pH for the FDDH calculations, that is absent for DH, and which is a sharp deviation from most proteins, giving a clear tendency towards maximal stability at acidic pH. The reducing stability as pH rises from acid pH is due to buried aspartic and glutamic acid sidechains that have elevated pKas, as noted previously for (non-human) pepsin (Lin *et al*., 1993). We demonstrate (Figure 7) that this variation is not removed by calculation with complete removal of basic sidechains (mutation of Lys and Arg to Met and His to Leu in 3utl), but is absent with the additional mutation (to Leu) of 5 buried acidic groups in 3utl (D96, E107, D149, D215, D303). Only one of these groups (D215) is part of the specific aspartic acid catalytic machinery, so most of the effect originates from outside of the active site. Second, and equally significant, is that even with the complete absence of basic sidechains, through modelled mutation, pepsin still exhibits a significant stabilisation of the folded form at pH 3 (Figure 7). This must arise from hydrogen-bond interactions to acidic groups, consistent with the exemplar graphical observations shown in Figure 6, and evidence for a systematic mechanism for using acidic groups to specifically stabilise protein folds in pH regions where stability is lost through neutralisation of salt-bridges.

**Figure 7:**
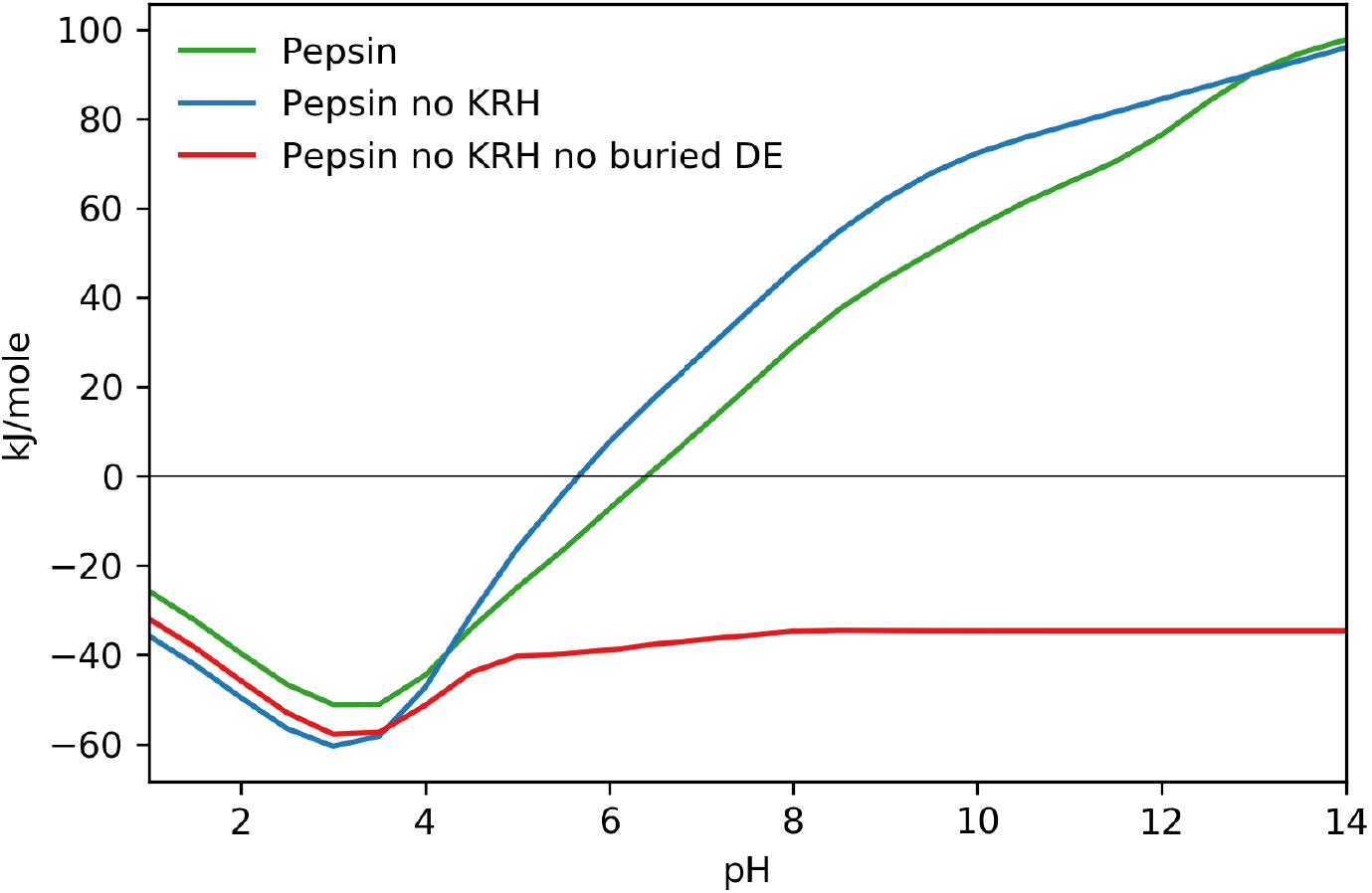
FDDH calculations of pH-dependence for human pepsin. Unaltered pepsin (green) is shown alongside pepsin with all basic amino acids mutated to non-polar amino acids blue), and with the further mutation of buried acidic amino acids to non-polar groups (red).

## Conclusions

The potential of therapeutic antibodies and next generation protein pharmaceuticals is well known, however issues remain with optimisation of production. In particular, the low pH hold required for viral inactivation and protein A chromatography leads to aggregation which can limit the development of therapeutic antibodies. We present a computational study of pH-dependence for IgG domains, the qualitative trends of which are in agreement with experimental data. The CH1 domain of the Fab domain exhibits similar charge-charge contributions to folded state stability at low pH as proteins that have evolved to reside in low pH environments. However, the similarity is limited, CH1 domains are not stable folded proteins in isolation, and thus mechanisms for designing in acid-stability are suggested that focus on proteins that have evolved to function at low pH. For the CH2 domain, the same groups that provide extensive stabilisation at neutral pH lead to destabilisation at acidic pH. This observation is consistent with experimental data showing that CH2 folded state stability is more acid sensitive than that of the CH3 domain, giving us confidence that the modelling can be used to consider engineering for acid stability. Comparison of proteins within the same family, but evolved to function at widely differing pH values leads to the suggestion that, in the low pH environment, basic sidechains (that are paired with acidic sidechains) can be swapped out for polar hydrogen-binding sidechains. Although ionisable group interactions are not the only stabilising force available to the protein, differences in stability at neutral pH and acidic pH are likely to be related to changes in the ionisation state of acidic and basic residues. A better understanding of these forces may provide insights for engineering acid-stable therapeutics in the future.

## Acknowledgements

The authors would like to thank members of the Warwicker group for feedback and support, and the computational shared facility at the university of Manchester.

