## Supplementary Figure 1 for "Modelling of pH-dependence to develop a strategy for stabilising mAbs at acidic steps in production"

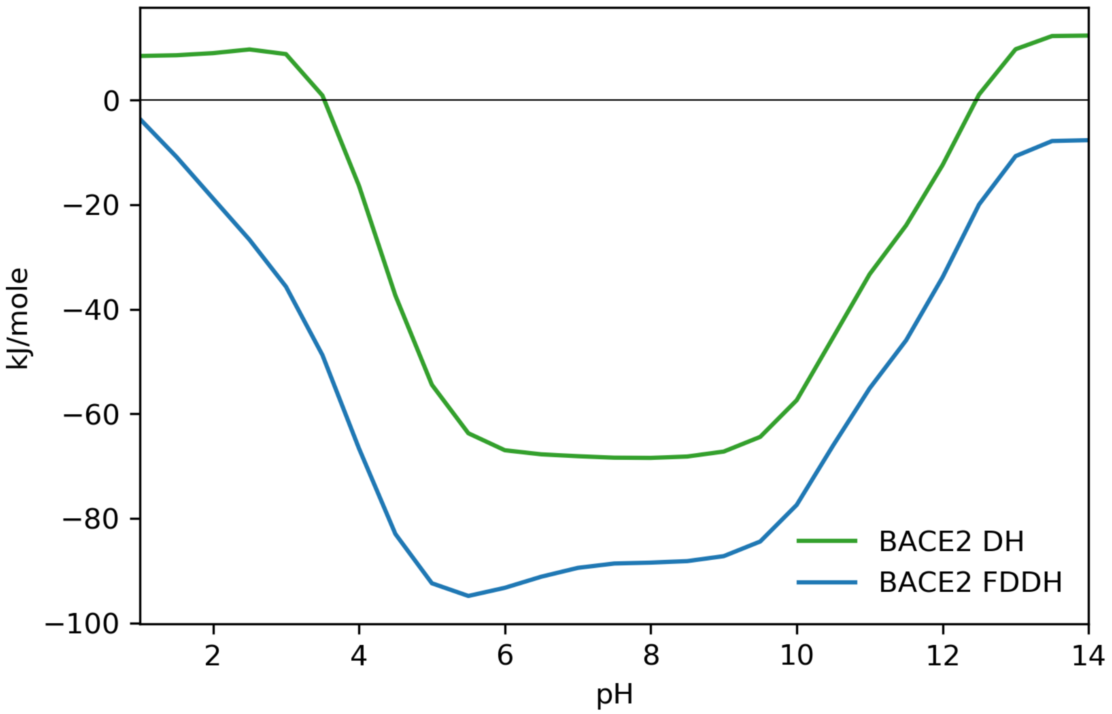


Figure S1: Comparison of acidic group interactions in equivalent regions of the homologues pepsin and BACE2.
